# A whole genomic CRISPR-Cas9 screen identifies the amino acid transporter *SLC43A1* (LAT3) as a major determinant of oxaliplatin sensitivity in colorectal cancer cells

**DOI:** 10.1101/2025.04.21.649594

**Authors:** NR Pawar, HM Wade, Z Jackson, N Poungpeth, AV Mitchell, CP Jewell, D Chan, RW Robey, PJ Batista, LM Jenkins, MM Gottesman

**Affiliations:** Laboratory of Cell Biology, Center for Cancer Research, National Cancer Institute, National Institutes of Health, Bethesda, MD

## Abstract

Colorectal cancer (CRC) is the second leading cause of cancer deaths in the United States, with a five-year survival rate of 65%. Oxaliplatin was the first platinum drug shown to improve CRC patient outcomes and is now a common adjuvant therapy for advanced disease, yet 90% of patients develop resistance. Oxaliplatin was developed as a third-generation derivative of cisplatin, but recent evidence points to divergent modes of action. Here, genome-wide CRISPR activation and knockout screens were conducted to identify genetic changes that confer resistance to oxaliplatin in two CRC cell lines with distinct molecular backgrounds (SW620 and RKO). Guide RNAs corresponding to the neutral amino acid transporter *SLC43A1* (LAT3) were the most significantly enriched in knockout screens and depleted in activation screens, suggesting a potential role for LAT3 in modulating oxaliplatin resistance. *In vitro* CRISPR knockout and overexpression of LAT3 in SW620 and RKO cell lines confirm increased resistance or sensitivity to oxaliplatin, respectively. Further analysis demonstrates that increased LAT3 levels corrrelate with increased intracellular levels of oxaliplatin, increased levels of DNA-platinum adducts and DNA damage, demonstrating that enhanced LAT3-mediated uptake of oxaliplatin can exert its expected mechanism of action and induce cytotoxicity. These findings may lead to a better understanding of oxaliplatin’s mode of action in CRC and can provide new insights into the interplay between essential nutrient uptake and drug transport.

**STATEMENT OF SIGNIFICANCE:** Oxaliplatin resistance remains a major clinical challenge for CRC treatment. Our study identified a novel role for LAT3 as a modulator of oxaliplatin sensitivity, offering new insights into drug resistance mechanisms.

## INTRODUCTION

Colorectal cancer (CRC) is the second-leading cause of cancer death for women and men in the United State, and is expected to cause 52,900 deaths in 2025 (1). The five-year survival rate for all stages of CRC is 65% but drops dramatically to 18% upon distant metastasis (2). CRC is generally considered a heterogenous disease, with mutations in *TP53* and *KRAS* (∼40-50% of cases (3,4)), and *BRAF* (∼10% of cases (5)) contributing to aggressive tumors that are difficult to treat. Surgery and localized radiation are the standard of care for early-stage disease, while adjuvant combination chemotherapy consisting of 5-fluorouracil (5-FU), leucovorin, and oxaliplatin (FOLFOX) is used for metastatic disease, generally resulting in a 50% response rate (6). Despite this favorable response and incremental advances in extending survival, many patients succumb to the disease after developing resistance to oxaliplatin-based adjuvant therapy.

Oxaliplatin, a third-generation platinum derivative, binds to DNA to form adducts, resulting in irreversible transcriptional errors and cellular apoptosis, thus impairing tumor cell division and growth (7,8). Although the precise mechanism of oxaliplatin cytotoxicity is largely inferred from that of the first-generation platinum cisplatin, it is thought that oxaliplatin requires fewer intra-strand links and DNA adducts for effective cytotoxicity (9), and may induce ribosomal biogenesis stress rather than DNA damage to kill cells, suggesting the possibility of other mechanisms of action (10). Oxaliplatin is reported to show stronger activity in colorectal and other gastrointestinal cancers while cisplatin and carboplatin are largely ineffective (11). Some studies show enhanced efficacy of oxaliplatin in conditions of cisplatin resistance, while others claim that the use of oxaliplatin is merely preferred due to reduced off-target toxicity, allowing synergy with other chemotherapeutic compounds (12). Commonly proposed mechanisms of platinum resistance thus far include enhanced drug detoxification, decreased uptake, metabolic rewiring, and alterations in DNA repair, cell death, or cell-signaling mechanisms (13). Despite some evidence of specific genes linked to chemoresistance, several may be activated simultaneously and can be difficult to target therapeutically. Understanding distinct molecular mechanisms underlying oxaliplatin-mediated anti-tumor properties and resistance is crucial for improving treatment strategies.

High-throughput, unbiased genetic screening using the CRISPR/Cas9 system is a comprehensive approach to investigate the molecular basis of multifactorial drug resistance in human cancers (14). Several studies using this technology in recent years have identified genes that predict drug response, mechanisms of action, and druggable targets upon treatment with a wide range of chemotherapeutics and targeted therapies (15). In this study, we perform genome-wide CRISPR/Cas9 knockout and activation screening in CRC cells with oxaliplatin treatments to systematically evaluate mechanisms of oxaliplatin resistance. We identified a major sodium-independent transporter of large neutral branched-chain amino acids, solute carrier family 43 member 1 *SLC43A1* (which encodes the protein L-type amino acid transporter 3, LAT3) among the genes whose expression is significantly altered under each screening condition. LAT3 is expressed in adult colon, small intestine, ovary, spleen, pancreas, liver, and skeletal muscle (16). It was initially found to be overexpressed in prostate cancer, and has since been associated with increased cellular uptake of leucine, leading to enhanced cell growth and proliferation (17). We observed that LAT3 levels in CRC cells correlated with sensitivity to oxaliplatin, uptake of oxaliplatin, and subsequent DNA damage caused by DNA-platinum adducts. Our data suggest that these LAT3 functions are specific to oxaliplatin compared to other platinum agents and were independent of its normal amino acid transport function. These data identify LAT3 as a novel determinant of oxaliplatin uptake in CRC cells, which affects sensitivity to this commonly used platinum agent.

## RESULTS

### Genome-wide CRISPR screens of CRC cell lines reveal *SLC43A1* (LAT3) as a top gene correlated with oxaliplatin sensitivity

To investigate key genes involved in oxaliplatin resistance in CRC, we conducted genome-wide CRISPR/Cas9 screens in two CRC lines (SW620 and RKO) with distinct molecular backgrounds (*KRAS*/*TP53* mutant; *BRAF* mutant/*TP53* WT, respectively). After generating cell lines with stable expression of Cas9 or dCas9-VP64, we utilized the human CRISPR Brunello (18) or CRISPR Calabrese (19) pooled libraries to conduct sgRNA-directed knockout (CRISPR-KO) or transcriptional activation (CRISPRa), respectively. Our goal was to investigate both positive and negative selection of sgRNAs after 20 days of oxaliplatin treatment to ensure a comprehensive and unbiased approach (**Fig. 1A**). Untreated cells were sampled at Day 0 and Day 20 for a time-matched control. In both knockout screens, we observed depletion of gRNAs targeting core essential genes in the T20 untreated condition compared to T0 control (**Fig. S1A**), confirming effective gene perturbation and capture of genes related to cell fitness under these screening conditions. In total, our CRISPRa screen identified the depletion of 156 and 222 genes in SW620 cells (under 0.38µM and 0.5µM selection, respectively), and 7491 genes in RKO cells associated with increased survival under oxaliplatin treatment (p<0.05, fold change ≥ 2, **Fig. S1B**). Our CRISPR KO screen identified the knockout of 301 and 446 genes in SW620 cells (under 0.38µM and 0.5µM selection, respectively) and 782 genes in RKO cells associated with increased survival under oxaliplatin treatment (p<0.05, fold change ≥ 2, **Fig. S1B)**. Upon gene ontology pathway analysis, sgRNAs targeting genes involved in DNA damage repair, transcription, translation and cell cycle transition/arrest were significantly depleted in all of the knockout screens (**Fig. S1C**), confirming our ability to capture known mechanisms of oxaliplatin resistance using this approach (13,20).

**Figure 1.**
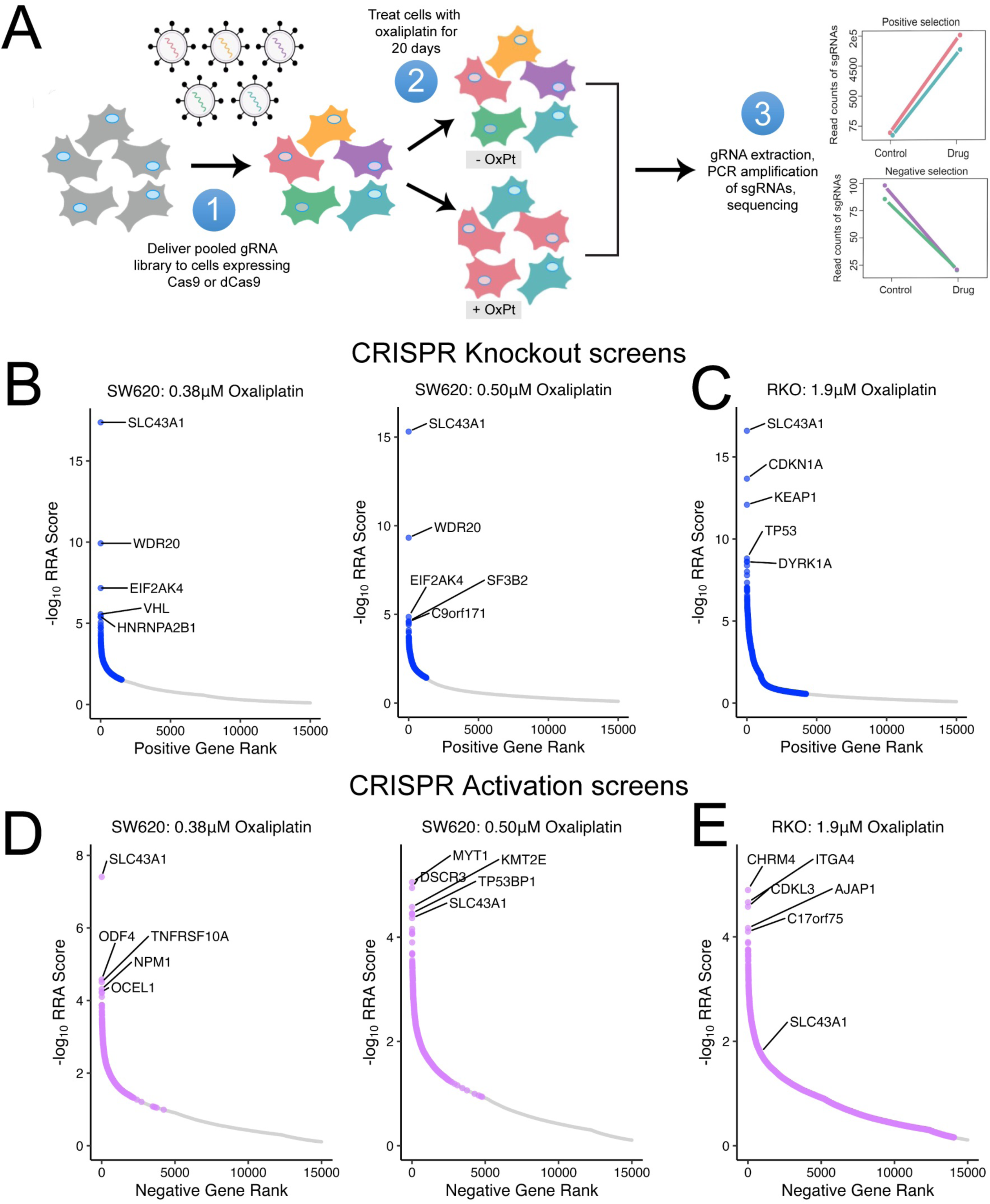
Genome-wide CRISPR screens of CRC cell lines reveal *SLC43A1* (LAT3) as top gene associated with oxaliplatin sensitivity. **A)** Overview of CRISPR screen methodology (adapted from (14)). **B)** SW620 KO screen;-log_10_ RRA scores in cells treated with 0.38µM (*left*) or 0.5µM OxPt (*right*) at Day 20 compared to control untreated cells at Day 20. **C)** RKO KO screen;-log_10_RRA scores in cells treated with 1.9µM OxPt at Day 20 compared to control untreated cells at Day 20. **D)** SW620 CRISPRa screen;-log_10_ RRA scores in cells treated with 0.38µM (*left*) or 0.5µM OxPt (*right*) at Day 20 compared to control untreated cells at Day 20. **E)** RKO CRISPRa screen;-log_10_ RRA scores in cells treated with 1.9µM at Day 20 compared to control untreated cells at Day 20. In each graph, top 5 hits are labeled; significant hits are colored (blue or lilac) while non-significant hits are in grey.

Notably, 5 positively selected hits were common across all knockout screen conditions, including *SLC43A1* (**Fig. S1B**). We found that upon treatment with two doses of oxaliplatin, SW620 cells had the most significant enrichment of sgRNAs targeting *SLC43A1* compared to untreated cells at Day 20 (**Fig. 1B**) and untreated cells at Day 0 (**Fig. S1D**). Similarly, *SLC43A1* was the most significantly enriched sgRNA target in the RKO knockout screen compared to either control (**Fig. 1C** and **Fig. S1E**). Importantly, sgRNAs targeting *SLC43A1* were not significantly altered between T20 and T0 untreated control conditions (**Fig. S1F**), suggesting the effects of SLC43A1 knockout are not simply due to general changes in cell fitness. Therefore, these data support SLC43A1 depletion as strong driver of survival under oxaliplatin treatment. In line with these findings, sgRNA-mediated gene activation resulted in reciprocal observations, whereby oxaliplatin selection resulted in a strong negative selection for sgRNAs targeting *SLC43A1*. We identified *SLC43A1* to be one of the most significant negatively selected CRISPRa hits in SW620 cells (ranked 1 of 1198 upon 0.38µM treatment; 5 of 1311 upon 0.5µM treatment; **Fig. 1D** and **Fig. S1G**). Similarly, RKO cells treated with oxaliplatin also exhibited a significant depletion of *SLC43A1*-targeted sgRNAs in the CRISPRa screen (**Fig. 1E** and **Fig. S1H**). Overall, *SLC43A1*-directed sgRNAs displayed a strong positive selection in the knockout screen (∼7 to10 log-fold in SW620 cells, ∼10 to12 log-fold in RKO cells) and a significant negative selection in the activation screen (-5 to-7 log fold in SW620,-8 to - 9 log fold change in RKO cells; **Fig. S1I**). These data suggest that depletion of *SLC43A1* gene expression confers resistance to oxaliplatin, while activation of *SLC43A1* expression confers increased sensitivity to oxaliplatin under our selection conditions.

### LAT3 knockout and overexpression in SW620 and RKO cells results in enhanced resistance and sensitivity to oxaliplatin, respectively

To validate the results from the CRISPR KO screen, we generated stable LAT3 knockout cell lines using single-gene CRISPR-KO and homology-directed repair plasmids. To validate the results from the CRISPRa screen, we generated stable LAT3 overexpression cell lines using a pcDNA3.1-SLC43A1 plasmid and matched empty vector controls. We confirmed efficient knockout and overexpression of LAT3 protein in SW620 cells by Western blot analysis in two individual clones each (**Fig. 2A**). LAT3 levels and plasma membrane localization were also confirmed by immunohistochemistry (IHC) staining (**Fig. 2B**). The effects of LAT3 knockout and overexpression on SW620 cells were measured by a luminescence-based Cell Titer-Glo cell viability assay upon treatment with oxaliplatin for 72 hours. We observed that both LAT3 knockout (LAT3KO) clones, B6 and B7, showed a clear increase in oxaliplatin resistance, as displayed by increased GI50 values (∼0.5µM versus ∼1µM; **Fig. 2C**). In contrast, we observed that both LAT3 overexpressing (LAT3OE) clones, C4 and C6, displayed a substantial increase in oxaliplatin sensitivity compared to empty vector controls, as displayed by decreased GI50 values (0.8µM versus ∼0.2µM; **Fig. 2D**). We also sought to validate the CRISPR KO and CRISPRa screen results in RKO cells using the same CRISPR-knockout and plasmid overexpression methods. LAT3 levels were depleted in two RKO knockout lines, LAT3KO-1 and-11, while three individual LAT3OE clones, 1, 5 and 9, displayed higher LAT3 protein levels compared to an empty vector control, as analyzed by Western blot (**Fig. 2E**) and IHC staining (**Fig. 2F**). Similar to the SW620 LAT3KO cell lines, both RKO LAT3KO clones displayed an increase in oxaliplatin resistance, as displayed by about a 4-5-fold increase in GI50 (**Fig. 2G**). RKO LAT3OE-1 and-9 cells also displayed a dramatic increase in oxaliplatin sensitivity, as shown by a ∼3-5-fold decrease in GI50 for LAT3OE-1 (**Fig. 2H**). It is worth noting that LAT3OE-9 cells had dramatic overexpression of LAT3 protein, correlating with the highest level of sensitivity to oxaliplatin (∼31-fold decrease in GI50). This demonstrates a correlation between LAT3 protein levels and oxaliplatin sensitivity. To further confirm changes in oxaliplatin sensitivity in these cell lines with a longer treatment period, we utilized colony formation assays. SW620 and RKO LAT3KO cells displayed a significantly higher survival fraction compared to respective parental cells (**Fig. 2I**). In comparison, SW620 and RKO LAT3OE cells exhibited a significantly lower survival fraction compared to pcDNA control cells (**Fig. 2J**), further confirming a role for LAT3 in mediating oxaliplatin sensitivity.

**Figure 2.**
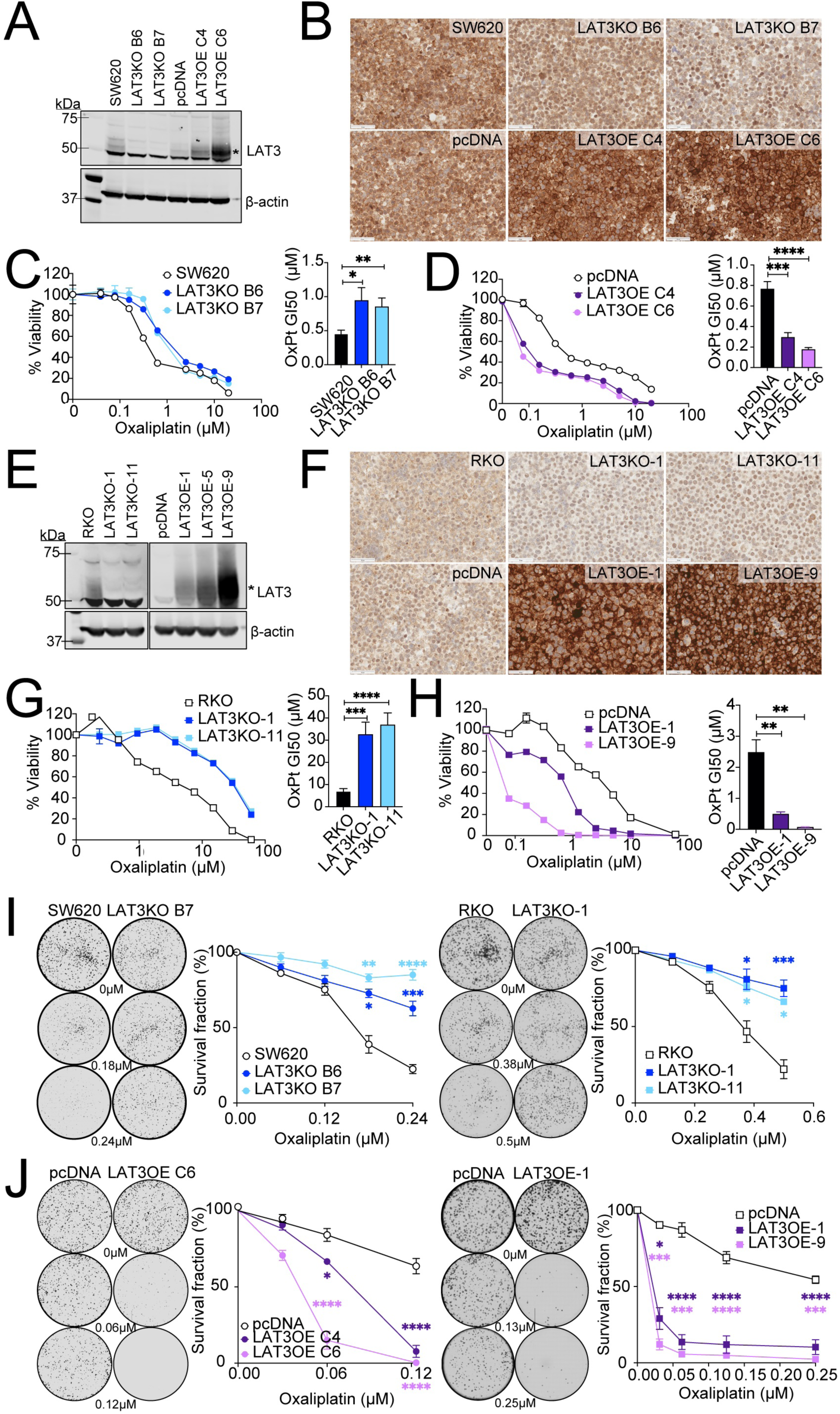
LAT3 knockout and overexpression in SW620 and RKO cells results in enhanced resistance and sensitivity to oxaliplatin, respectively. **A)** Immunoblot analysis of SW620 parental cells, LAT3KO clones B6 and B7, pcDNA vector control, and LAT3OE clones C4 and C6 for LAT3 and β-actin protein expression. Asterisk indicates band corresponding to LAT3, above nonspecific band ∼50kDa. **B)** Cell pellets were analyzed using immunohistochemistry for LAT3 expression. Representative images were taken at 40X magnification; scale bars represent 50µM. **C)** SW620-LAT3KO and **D)** LAT3OE cells were treated with oxaliplatin (0-20µM) for 72 hours and cell viability was measured by CellTiterGlo luminescence-based assay. Representative dose response curves show average cell viability from triplicate wells; bar graphs show an average GI50. **E)** Immunoblot analysis of RKO parental cells, LAT3KO clones 1 and 11, pcDNA vector control, and LAT3OE clones 1, 5 and 9 for LAT3 and β-actin protein expression. **F)** Cell pellets were analyzed using immunohistochemistry for LAT3 expression. Representative images were taken at 40X magnification; scale bars represent 50µM. **G)** RKO-LAT3KO and **H)** LAT3OE cells were treated with oxaliplatin (0-60µM) for 72 hours and cell viability was measured by CellTiterGlo. Representative dose response curves show average cell viability from triplicate wells; bar graphs show average GI50. **I)** SW620 and RKO LAT3KO or **J)** LAT3OE cells were seeded in 6-well plates at 1,000 cells/well, treated with 0-0.5µM oxaliplatin for 10-12 days, and colonies were stained with crystal violet. Representative whole-well scans show stained colonies at experimental endpoint. Graphs represent average survival fraction from 3 independent experiments performed in triplicate. All error bars represent SEM. Data are represented as mean ± SEM; n = 3 independent experiments; *p<0.05, **p<0.01, ***p<0.005, ****p<0.001.

### LAT3 does not mediate sensitivity to other platinum chemotherapeutic agents in SW620 and RKO cells

Oxaliplatin is generally thought to adopt a similar mechanism of action to previously derived platinum agents such as cisplatin and carboplatin. However, evidence of diverging pathways and context-dependent instances of cross-resistance in cancer types have been reported. To investigate whether LAT3 may play a role in sensitivity to other platinum-based anti-cancer agents, we treated SW620 and RKO LAT3KO and LAT3OE cells with cisplatin, carboplatin, or phenanthriplatin, and measured cell viability after 72 hours. Except for a small increase in sensitivity to carboplatin in the overexpressing cells, neither LAT3 knockout (**Fig. S2A-B**) nor LAT3 overexpression (**Fig. S2C-D**) in either cell line significantly affected sensitivity to any of these platinum-based chemotherapeutic agents to the same degree as oxaliplatin, suggesting a specific role of LAT3-mediated cytotoxicity in response to oxaliplatin in these CRC cells.

### LAT1 does not mediate sensitivity to oxaliplatin

LAT3 belongs to a family of four L-type amino acid transporters, including *SLC7A5* (LAT1), which is primarily responsible for transporting large neutral amino acids, particularly leucine (21). In our CRISPR KO screen using SW620 cells, sgRNAs targeting *SLC7A5* were significantly depleted in oxaliplatin-treated cells compared to untreated controls (**Fig. S3A**), leading us to hypothesize its possible opposite role to LAT3. To investigate the role of LAT1 in SW620 cells, we generated LAT1 CRISPR KO clones (1 and 2), and a LAT1 overexpressing clone (C9) and validated protein expression by Western blot (**Fig. S3B**). Treatment of LAT1KO clones (**Fig. S3C**) and LAT1OE cells (**Fig. S3D**) with oxaliplatin for 72 hours resulted in no significant difference in cell viability compared to parental or empty vector control cells. LAT1KO also did not affect survival fraction in a colony formation assay compared to parental cells (**Fig. S3E**), confirming that LAT1 does not have a role in mediating oxaliplatin resistance in these CRC cells. Protein and mRNA levels of LAT1 and other LAT family members LAT2 (*SLC7A8*), or LAT4 (*SLC43A2*) are not altered under LAT3 knockout or overexpression conditions (**Fig. S3G-H**), indicating that none of these other neutral amino acid transporters are contributing to the oxaliplatin resistance phenotype.

### LAT3 determines intracellular accumulation of oxaliplatin in CRC cells

LAT3 has been demonstrated to import large neutral branched-chain amino acids (22). We hypothesized that LAT3 mediates oxaliplatin sensitivity by determining intracellular accumulation of oxaliplatin in CRC cells. After treatment with 1µM oxaliplatin for three and six hours, LAT3KO cells displayed reduced levels of intracellular platinum compared to parental cells (**Fig. 3A**), while LAT3OE cells displayed significantly increased levels of intracellular platinum compared to empty vector controls (**Fig. 3B**), as measured by inductively coupled plasma mass spectrometry. These data support the hypothesis that LAT3 may directly or indirectly result in the transport of oxaliplatin into the cell to mediate cytotoxic effects. Upon similar treatment with cisplatin, there was no significant difference in intracellular levels of platinum in LAT3KO or LAT3OE cells compared to the respective controls (**Fig. 3C-D**), indicating that LAT3-determined transport is specific to oxaliplatin. LAT1KO cells did not display a significant difference in oxaliplatin uptake compared to parental SW620 cells (**Fig. S3F**), showing that this transport activity is specific to LAT3. The increased cellular platinum accumulation in LAT3OE cells was paralleled by an increase of total platinum bound to genomic DNA (**Fig. 3E**) and increased levels of phosphorylated γ-H2AX, a measure of DNA damage (**Fig. 3F**). In contrast, LAT3KO cells showed decreased DNA-Pt adducts and phosphorylated γ-H2AX levels (**Fig. 3E-F**), confirming proper localization of oxaliplatin upon uptake and resulting DNA damage to mediate cytotoxicity.

**Figure 3.**
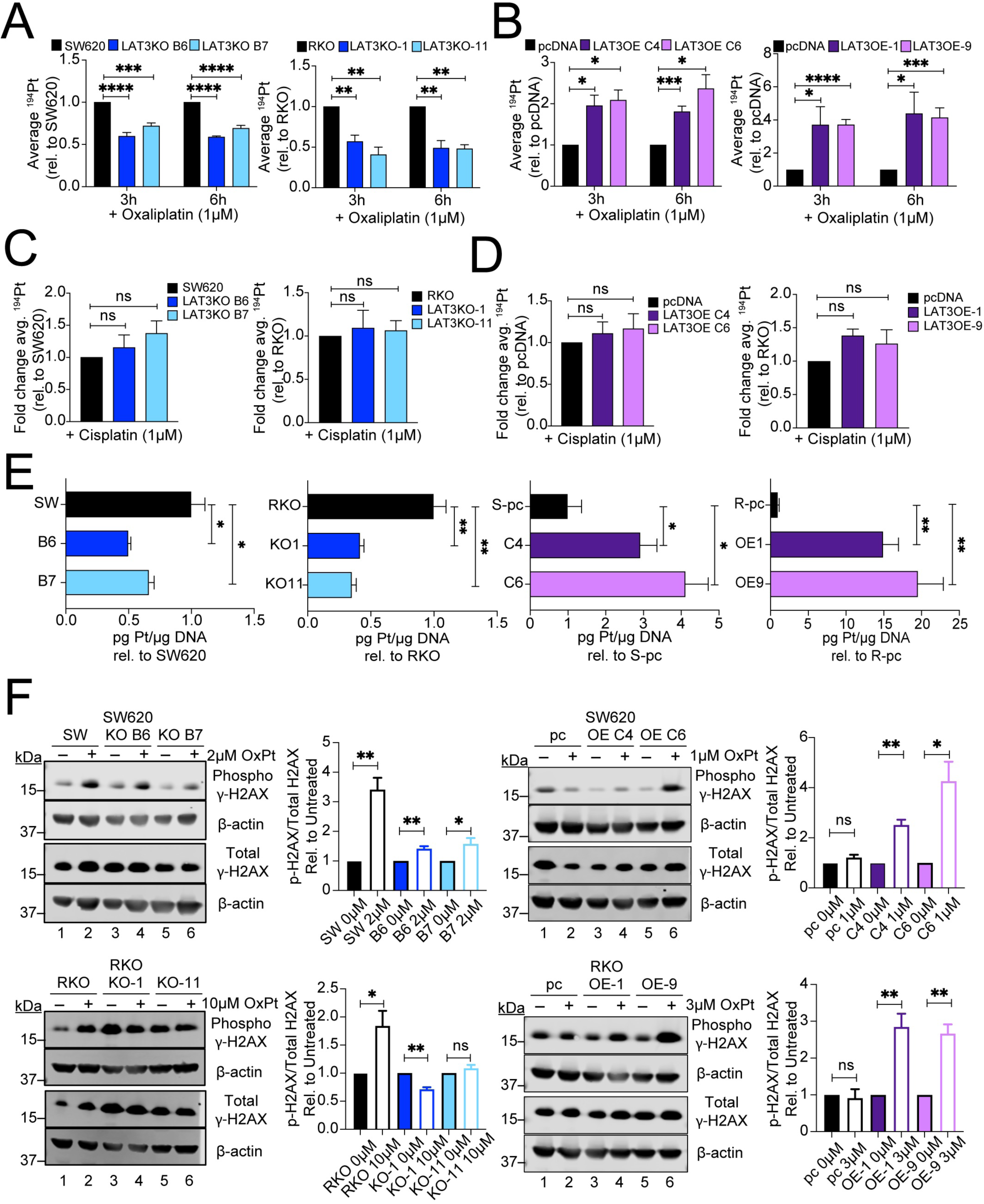
**LAT3 knockout in CRC cells reduces intracellular accumulation of oxaliplatin**. **A)** SW620 and RKO LAT3KO or **B)** LAT3OE cells were treated with 1µM oxaliplatin in complete growth media for 3 or 6 hours, washed with dPBS, and cell pellets were collected in dPBS to measure intracellular platinum accumulation via ICP-MS. Average ^194^Pt values were normalized to cell number, and data represents average fold change relative to parental SW620/RKO or pcDNA control cells. **C)** SW620 and RKO LAT3KO or **D)** LAT3OE cells were treated with 1µM cisplatin for 6 hours, washed, and cell pellets were collected to measure intracellular platinum accumulation via ICP-MS. Average ^194^Pt values were normalized to cell number, and data represents average fold change relative to parental SW620/RKO or pcDNA control cells. **E)** DNA was collected from cells treated with oxaliplatin (2µM, 10µM, 1µM, and 3µM respectively) for 24 hours; platinum adducts were measured by ICP-MS. Data represents average ^194^Pt measured in treated cells relative to respective controls. **F)** Cells were treated with oxaliplatin (as noted) and protein extracted from whole cell lysates after 24 hours. γ-H2AX protein levels were measured by western blot (representative images shown). Phosphorylated γ-H2AX and total γ-H2AX levels were measured by densitometric analysis on ImageJ. Each was normalized to β-actin levels and are shown as average fold change compared to respective untreated controls. All error bars represent SEM. Data are represented as mean ± SEM; n = 3 independent experiments performed with 3-4 replicates; *p<0.05, **p<0.01, ***p<0.005, ****p<0.001, ns = not significant.

### LAT3-mediated sensitivity to oxaliplatin is independent of amino acid transport

Since LAT3 is an amino acid transporter, we hypothesized that oxaliplatin may rely or “piggy-back” on leucine or other neutral amino acids to be transported by LAT3. The presence of excess amino acids in the extracellular environment does not enhance or restore influx of oxaliplatin in LAT3KO cells compared to parental cells (**Fig. 4A**), nor does it restore sensitivity to oxaliplatin (**Fig. 4B**). Conversely, depletion of extracellular amino acids does not prevent the enhanced oxaliplatin uptake mediated by LAT3 overexpression (**Fig. 4C**). Inhibition of LAT3’s leucine transport function by pan-LAT inhibitor BCH (23,24) also does not prevent the enhanced sensitivity mediated by LAT3 overexpression (**Fig. 4D**). These results suggest that LAT3 may mediate transport of oxaliplatin into cells independent of its amino acid transport functions.

**Figure 4.**
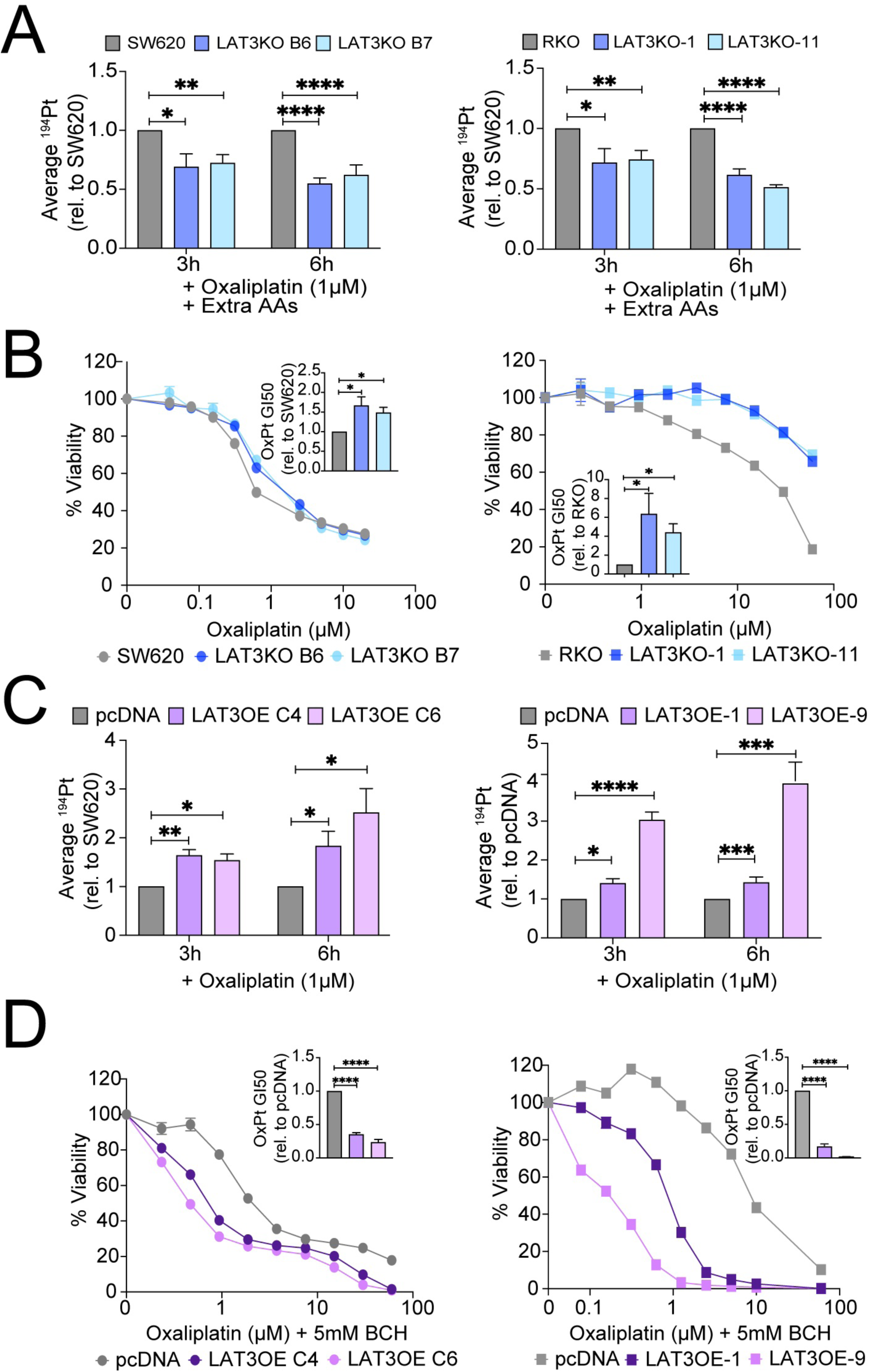
LAT3-mediated sensitivity to OxPt is independent of amino acid transport. **A)** SW620 and RKO LAT3KO cells were treated with 1µM oxaliplatin in complete growth media supplemented with extra amino acids for 3 or 6 hours, washed, and cell pellets were collected to measure intracellular platinum accumulation via ICP-MS. Average ^194^Pt values were normalized to cell number, and data represents average fold change relative to parental SW620 or RKO cells. **B)** SW620 and RKO LAT3KO cells were treated with 0-20µM or 0-60µM oxaliplatin, respectively, in the presence of extra amino acids for 72 hours and cell viability measured by CellTiterGlo. Representative dose response curves show average cell viability from triplicate wells; bar graph insets represent average GI50 values relative to parental controls. **C)** SW620 and RKO LAT3OE cells were treated with 1µM oxaliplatin in PBS (with Ca^2+^, Mg^2+^) for 3 or 6 hours, washed, and cell pellets were collected to measure intracellular platinum accumulation via ICP-MS. Average ^194^Pt values were normalized to cell number, and data represents average fold change relative to respective control pcDNA cells. **D)** SW620 and RKO LAT3OE cells were treated with 0-20µM or 0-60µM oxaliplatin, respectively, in the presence of pan-LAT inhibitor BCH (5mM) for 72 hours and cell viability measured by CellTiterGlo. Representative dose response curves show average cell viability from triplicate wells; bar graph insets represent average GI50 values relative to pcDNA controls. All error bars represent SEM. Data are represented as mean ± SEM; n = 3 independent experiments performed in triplicate; *p<0.05, **p<0.01, ***p<0.005, ****p<0.001.

### LAT3 partially mediates sensitivity to oxaliplatin in ovarian cancer cells

To investigate whether LAT3-mediated sensitivity is generalizable to other cancer types, we turned to ovarian cancer cell lines. Platinum-based chemotherapy is a crucial component of the standard of care regimen for ovarian cancer patients, the majority of which develop resistance (25). To determine if LAT3 expression plays a role in oxaliplatin resistance in ovarian cancer, we examined a panel of 1A9 ovarian cancer cells (derived from A2780 cells) that were selected with increasing concentrations of oxaliplatin ((26); **Fig. 5A**). Upon immunoblot analysis, we observed that loss of LAT3 expression was associated with increased oxaliplatin resistance (**Fig. 5B**), leading us to hypothesize that LAT3 may play a role in oxaliplatin resistance in ovarian cancer as well. Decreasing levels of LAT3 and increasing resistance also correlated with decreased intracellular accumulation of oxaliplatin (**Fig. 5C**), mirroring our observations in CRC cells. Independent 1A9-LAT3KO clones were generated by CRISPR/Cas9, and protein levels were confirmed by Western blot (**Fig. 5D**). Both 1A9 LAT3KO clones displayed an increase in resistance to oxaliplatin (**Fig. 5E**), whereas sensitivity to cisplatin and carboplatin remained unchanged (**Fig. 5F, Fig. S4A**). To investigate whether LAT3 expression would restore sensitivity to oxaliplatin resistant lines, we overexpressed LAT3 in 1A9-OX80 cell lines and confirmed expression by Western blot (**Fig. 5G**). LAT3OE #4 cells displayed partially decreased resistance to oxaliplatin compared to the OX80 resistant line (**Fig. 5H**). LAT3OE #4 cells also had a slight decrease in resistance to cisplatin, but showed no difference in sensitivity to carboplatin compared to OX80 cells (**Fig. 5I, Fig. S4B**). Notably, the effect on oxaliplatin sensitivity upon LAT3 perturbation in 1A9 cells was minimal compared to the more dramatic effect in CRC cells, indicating the likelihood of other resistance mechanisms in the 1A9 highly selected oxaliplatin-resistant ovarian cancer cells. 1A9-OxR cells were also cross-resistant to both cisplatin and carboplatin (**Fig. S4C**), and this corresponded with decreased uptake of cisplatin and carboplatin (**Fig. S4D**), further demonstrating that the role of LAT3 in the divergent modes of action for oxaliplatin may be cancer-type specific.

**Figure 5.**
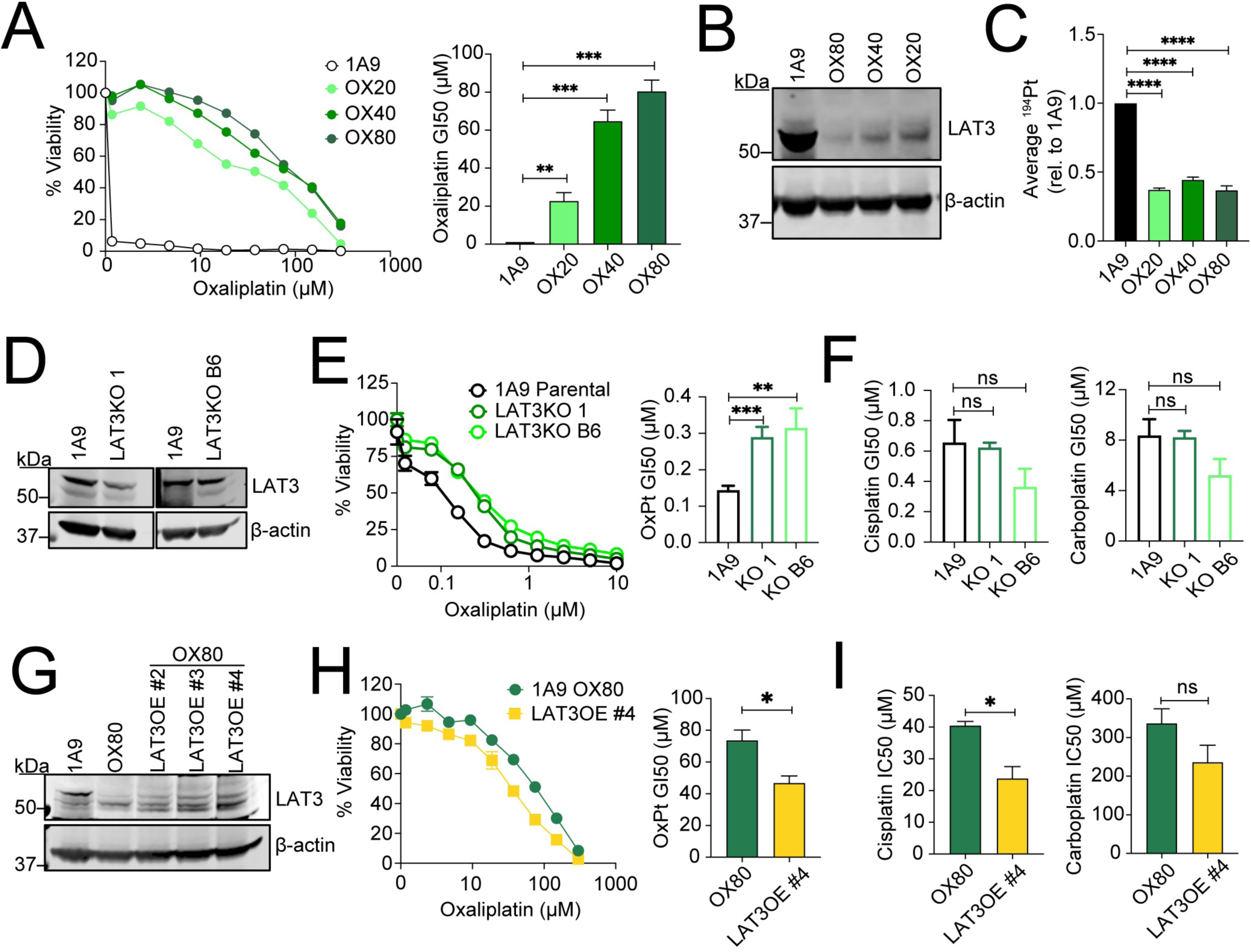
LAT3 partially mediates sensitivity to OxPt in 1A9 ovarian cancer cells. **A)** 1A9 ovarian cancer cells were exposed to 20, 40, and 80µM of oxaliplatin as described to generate resistant lines (26). Cells were treated with 0-300µM oxaliplatin for 72 hours and cell viability was measured by CellTiterGlo. Representative dose response curves show average cell viability from triplicate wells; bar graphs show average GI50. **B)** Immunoblot analysis of parental 1A9 and oxaliplatin resistant cell lines (OX80, OX20, and OX20) for LAT3 and β-actin protein expression. **C)** 1A9 parental and oxaliplatin resistant cells (cultured in the absence of oxaliplatin) were treated with 1µM oxaliplatin in complete growth media for 6 hours, washed with dPBS, and cell pellets were collected in dPBS to measure intracellular platinum accumulation via ICP-MS. Average ^194^Pt values were normalized to cell number, and data represents average fold change relative to parental 1A9 cells. **D)** Immunoblot analysis of parental 1A9 cells and two independent LAT3KO clones (LAT3KO 1 and LAT3KO B6) for LAT3 and β-actin protein expression. **E)** Cells were treated with 0-10µM oxaliplatin for 72 hours and cell viability was measured by CellTiterGlo. A representative dose response graph shows average cell viability from triplicate wells; bar graphs show average GI50s. **F)** Cells were treated with 0-10µM cisplatin or 0-100µM carboplatin for 72 hours and cell viability measured by CellTiterGlo; data represent average GI50s. **G)** Immunoblot analysis of parental 1A9 cells, 1A9-OX80 cells, and independent clones of 1A9-OX80 cells overexpressing LAT3 (#2, #3, and #4) for LAT3 and β-actin protein expression. **H)** Cells were treated with 0-300µM oxaliplatin and cell viability was measured by CellTiterGlo. A representative dose response graph shows average cell viability from triplicate wells; bar graph shows average oxaliplatin GI50. **I)** Cells were treated with cisplatin, and carboplatin for 72 hours, data represent average GI50s. All error bars represent SEM. Data are represented as mean ± SEM; n = 3 independent experiments performed in triplicate; *p<0.05, **p<0.01, ***p<0.005, ****p<0.001, ns = not significant.

### LAT3 levels mediate response to oxaliplatin treatment in CRC

To investigate a clinically relevant role for LAT3 in CRC, we first analyzed LAT3 mRNA levels in tumor and normal patient samples across tumor types using publicly available RNASeq data (TNMPlot pan-cancer analysis; https://tnmplot.com/analysis (27)). Most notably, levels of LAT3 in colon and rectal tumor samples were elevated compared to normal tissues (**Fig. 6A**). Furthermore, we analyzed LAT3 mRNA levels in cell lines across tumor types and of CRC origin, stratified by their response to oxaliplatin treatment *in vitro*, using publicly available transcriptomic data (ROC Plotter; https://rocplot.org/cells (28)). LAT3 levels were elevated in tumor lines that were more sensitive to oxaliplatin treatment (**Fig. 6B**), further indicating a role for LAT3 in regulating sensitivity to oxaliplatin.

**Figure 6.**
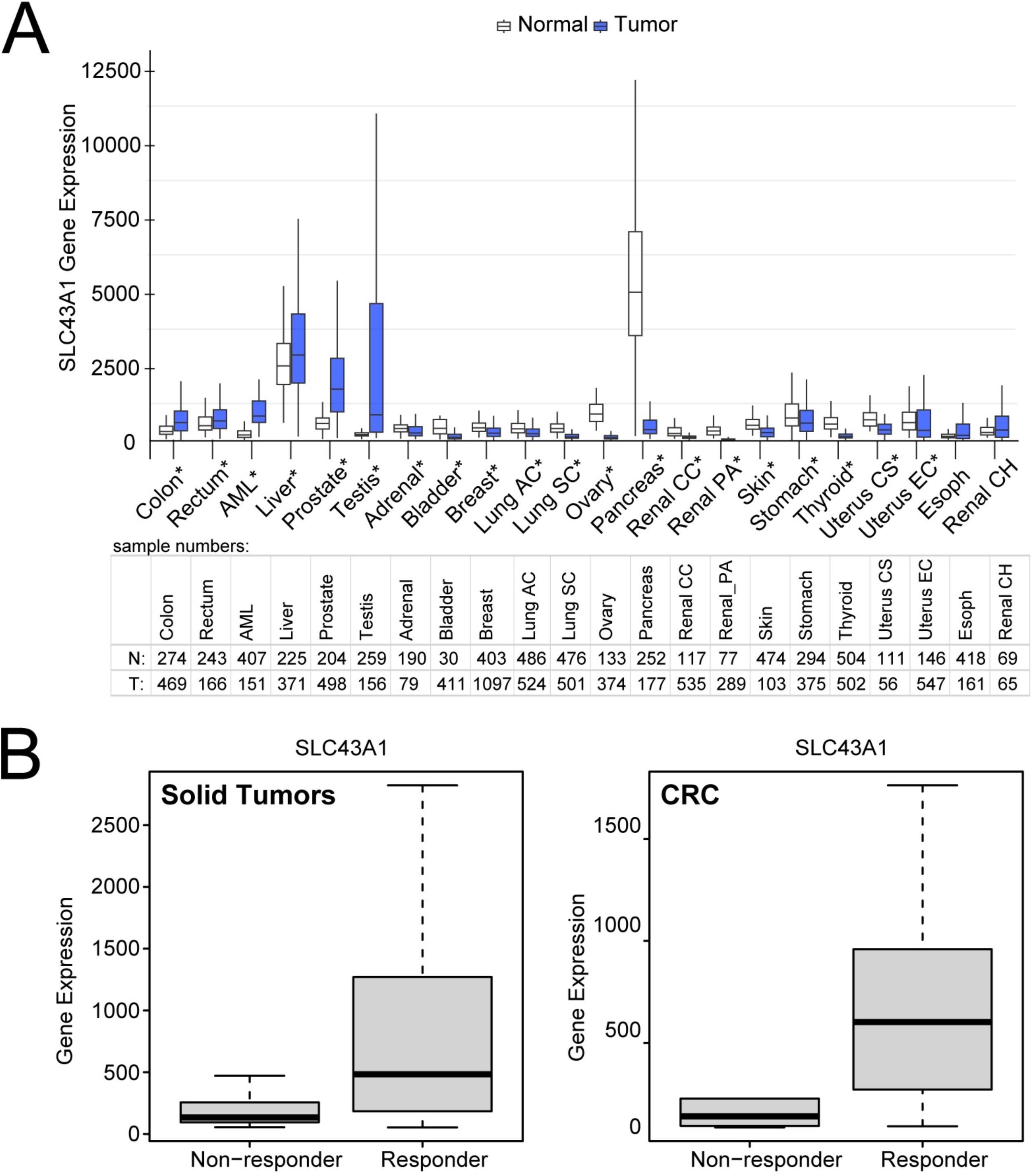
LAT3 is upregulated in CRC patient tumors and can be a determinant of oxaliplatin sensitivity *in vitro*. **A)** SLC43A1 RNA levels were determined in normal and cancer tissues using the TNMPlot pan-cancer analysis (https://tnmplot.com/analysis/). Asterisks indicate Mann-Whitney p < 0.05 and expression >10 in tumor or normal tissues. Sample numbers for normal (N) and tumor (T) are listed in table below (Lung AC = Lung adenocarcinoma; Lung SC = Lung squamous cell carcinoma; Renal CC = Renal clear cell carcinoma; Renal PA = Renal papillary cell carcinoma; Uterus CS = Uterine carcinosarcoma; Uterus EC = Uterine corpus endometrial carcinoma; Renal CH = Renal chromophobe). **B)** RNA levels of SLC43A1 were determined in cell lines from all solid tumors (*left*) and colon/colorectal cancer (*right*) and stratified by response to oxaliplatin based on upper and lower tertiles of GI50. Data from the GDSC2 datasets were plotted on www.rocplot.org/cells. Solid tumor cell lines: responder N = 267, non-responder N = 265; ROC p-value = 0, Mann-Whitney test p-value = 1e-26, fold change = 3. Colon/colorectal cancer cell lines: responder N = 26, non-responder N = 10; ROC p-value = 1e-08; Mann-Whitney test p-value = 9e-04, fold change = 3.6.

## DISCUSSION

Oxaliplatin remains a major component of the standard of care regimen for advanced CRC due to its potent activity as a platinum agent, despite the development of resistance, which ultimately results in poor patient outcomes. Our study was designed to investigate the genetic basis of oxaliplatin resistance using genome-wide CRISPR knockout and activation screens in two distinct *in vitro* models of CRC. Our findings identify and characterize a novel mechanism of oxaliplatin uptake mediated by LAT3 in CRC cells, which appears to be independent of its normal amino acid transport function and can regulate sensitivity to oxaliplatin by accumulation of DNA-Pt adducts and DNA damage.

Although initially derived from cisplatin to reduce off-target toxicity, evidence of both overlapping and differing modes of action have been reported for oxaliplatin. Reduced intracellular accumulation has been strongly correlated with increased resistance for both cisplatin and oxaliplatin in many cancer cell lines (29), although detailed mechanisms for oxaliplatin are as yet uncharacterized. One major way the oxaliplatin mechanism of action is thought to diverge from that of cisplatin is its relatively higher reliance on active transport versus passive diffusion into cells, especially at longer time points (30). In some cells, uptake of oxaliplatin can be facilitated by the human copper transporter hCTR1, organic cation transporters OCT1, 2, and 3 (*SLC22A1-3*) and multidrug and toxin extrusion MATE1 transporter (30). Decreased expression of *SLC22A2* (OCT2) has been reported to be associated with decreased cytotoxicity to oxaliplatin *in vitro* (31); however, this transporter is primarily expressed in renal cells and in dorsal root ganglia, suggesting a role in peripheral neurotoxicity as opposed to uptake by colorectal cancer cells (32). Given the contribution of active transporters to oxaliplatin accumulation in cells, it is perhaps not surprising that our genome-wide CRISPR screens identified a new transporter in CRC cells as shown by a dramatic alteration in sgRNAs directed towards the neutral amino acid transporter LAT3. OCT2 mRNA expression was not detected in SW620 or RKO cells (data not shown). In fact, these previously proposed transporters of oxaliplatin were not significantly altered under our CRISPR/Cas9 screening conditions (**Fig. S5A**) and had inconsistent variation upon LAT3 manipulation in LAT3KO and LAT3OE cells (**Fig S5B**), leading us to conclude that the regulation of platinum accumulation to mediate resistance is cancer-type specific.

Our results indicate that CRC cells may downregulate LAT3 as a mechanism of resistance to reduce oxaliplatin accumulation in the cell. Previous studies have suggested that LAT3 can be transcriptionally regulated by androgen receptor signalling (23,33) or EGF-activated PI3K/Akt signalling (34) in prostate cancer, while others report regulation of LAT3 by oncogenic MYC expression (35), but these mechanisms have yet to be studied in the context of CRC and response to oxaliplatin treatment. CRC patient tumors, which are initially oxaliplatin sensitive, display increased LAT3 mRNA levels compared to normal tissue, while many other tumor types tend to downregulate LAT3 compared to normal tissue (i.e. breast, lung, ovary, pancreas; **Fig. 6A**). This upregulation of the LAT3 transporter in CRC may explain the enhanced efficacy of oxaliplatin in CRC compared to other cancer types. Our data also showed a consistent role for LAT3 across mutational backgrounds common in patients, suggesting it can be a useful prognostic tool to combat heterogenous and aggressive CRC. There is little to no exploration of oxaliplatin being used as a therapeutic agent for AML, liver (36), prostate (37), and testicular (38) cancers, and perhaps the upregulation of LAT3 observed in these tumor tissues could facilitate an initial response. *In vitro* studies indicate that cell lines across tumor types and CRC cell lines that are responsive to oxaliplatin have increased levels of LAT3 compared to those that are more resistant, further suggesting a LAT3-mediated mechanism for oxaliplatin resistance. Additional studies investigating LAT3 in other downstream mechanisms of resistance, such as drug sequestration or inactivation, increased DNA repair, reduced apoptosis, or triggering nucleolar stress are necessary to better characterize this pathway.

Previous characterizations of acquired platinum resistance in cell lines selected over long periods of time propose a pleiotropic response (39), and any changes in overall accumulation could require simultaneous inactivation of many of the suggested transporters (29). A benefit of our CRISPR/Cas9 approach is the ability to perturb one gene per cell and observe resulting phenotypic differences under drug selection, allowing a characterization of individual genes involved in resistance. While the CRISPR/Cas9 screening approach to investigate multidrug resistance has resulted in intriguing discoveries, some limitations remain. Our study focuses on molecular and phenotypic changes after single-agent treatment, while currently oxaliplatin is only FDA-approved for usage as one component of the FOLFOX regimen to treat advanced CRC. This combination treatment also limits our interpretation of patient samples before and after treatment. This study also does not account for the effect of the tumor microenvironment or three-dimensional cell-cell interactions that play a crucial role in *in vivo* drug uptake and resistance. Further validation using a physiologically relevant model such as tumor organoids, and clinically achievable regimens and dosages, may further enhance the biological significance of our findings. Still, it is intriguing to speculate that a primary determinant of the response of CRC to oxaliplatin treatment might relate to the expression and function of LAT3 in these cells.

One unresolved issue is the mechanism by which LAT3 controls the accumulation of oxaliplatin in cells. There are two plausible hypotheses: (1) LAT3 is itself the transporter and the oxaliplatin “piggy-backs” on this transport system at a site different from the neutral amino acid transport site; and (2) LAT3 indirectly controls the uptake of oxaliplatin through as yet unknown mechanism(s). At this time, we do not have any data to rule in or out either of these hypotheses.

Patients with tumors that have responded to platinum-based therapies were reported to have increased platinum accumulation compared to non-responders (40), signifying the importance of drug accumulation for clinical efficacy. Detailed clinical studies on regulators of oxaliplatin accumulation are lacking, but a better understanding can lead to personalized alterations of treatment (41) and prognostic biomarkers of resistance. There are reports of cisplatin nanocapsules that primarily enter cells by endocytosis and can bypass active influx mechanisms (42), suggesting an alternative mode to develop for oxaliplatin delivery in patients with resistant disease and decreased LAT3 levels.

## ACKNOWLEDGEMENTS

This work was supported by the Intramural Research Program of the NIH. We would like to thank members of the Molecular Histopathology Lab at NCI Frederick, Donna Butcher, Elijah Edmondson, and Stephanie Smith, for their assistance with IHC staining. We appreciate the helpful discussions with James P. Madigan. The authors also thank the Center for Cancer Research (CCR) Genomics Core in Bethesda, Maryland. This work utilized the computational resources of the NIH HPC Biowulf cluster (http://hpc.nih.gov).

## METHODS

### Chemicals

Oxaliplatin was obtained from MedChemExpress (Monmouth Junction, NJ, cat# HY-17371) and Sigma (St. Louis, MO, cat# O9512). Cisplatin (cat# P4394), carboplatin (cat# C2538) and phenanthriplatin (cat# SML2811) were obtained from Sigma. BCH (pan-LAT inhibitor) was obtained from Cayman Chemical (Ann Arbor, MI, cat# 15249).

### Cell culture

SW620 (colorectal) cells were obtained from the Division of Cancer Treatment and Diagnosis Tumor Repository, National Cancer Institute at Frederick, MD. RKO (colorectal) cells were purchased from ATCC (Manassas VA). 1A9 cells (a derivative of A2780 ovarican cancer cells) and the oxaliplatin resistant sublines 1A9 OX20, OX40, and OX80 were a kind gift from Dr. Tito Fojo (Columbia University, New York, NY). SW620 and 1A9 cell lines were cultured in RPMI Medium + 10% FBS, 1% Pen/Strep, and 1% Glutamine. RKO (colorectal) cells were cultured in EMEM Medium + 10% FBS, 1% Pen/Strep, and 1% Glutamine. SW620 and RKO cells were authenticated by ATCC. All cells were grown at 37°C with 5% CO_2_ in a humidified environment.

### CRISPR-Cas9 Workflow

SW620 and RKO cell lines were transduced with Cas9 (lentiCas9-Blast was a gift from Feng Zhang (obtained from Addgene, Watertown, MA, viral prep #529622-LV; http://n2t.net/addgene:52962; RRID:Addgene_52962) (43)) or dCas9-VP64 (lentidCAS-VP64_Blast was a gift from Feng Zhang (Addgene plasmid # 61425-LVC; http://n2t.net/addgene:61425; RRID:Addgene_61425 (44)). The stable expression of Cas9 and dCas9 were confirmed by Western blotting. Cells were transduced with either the Brunello and Calabrese libraries, at a low MOI (∼0.3) to ensure effective barcoding of individual cells. The Human Brunello CRISPR knockout pooled library was a gift from David Root and John Doench (Addgene viral prep #73178-LV) (18). The Human Calabrese CRISPR Activation Pooled Library Set A was a gift from David Root and John Doench (Addgene viral prep #92379-LVC) (19). The transduced cells were selected with 600 ng/mL (RKO) or 2.3 μg/mL (SW620) of puromycin for 3 days to generate a cell pool carrying the libraries, which was then treated with oxaliplatin (1.9 μM for RKO cells or 0.38 μM and 0.5 μM for SW620 cells) for 20 days. After treatment, at least 1 × 10^9^ cells were collected to ensure over 500X coverage of the library. Genomic DNA was extracted using the QIAmp DNA blood Cell Maxi Kit (Qiagen, Germantown, MD) according to the manufacturer’s protocol. The sgRNA sequences were amplified using Taq polymerase (Takara Bio, Inc., San Jose, CA) and adapted for sequencing. The desired DNA product was purified with 6% TBE gel (Invitrogen, Waltham MA) and subjected to massive parallel amplicon sequencing carried out by an Illumina sequencer (Sanger/Illumina 1.9). The sgRNA read count and hits calling were analyzed by the MAGeCK v0.5.7 algorithm18. To assess the function of the negatively selected hits from the CRISPR KO screen, Gene Ontology (GO) Biological Process functional enrichment analysis was performed using the R package clusterProfiler (v 4.12.6 (45)).The top 500 most significantly depleted genes (p ≤ 0.05) from each CRISPR knockout screen were used as input for the analysis. Enrichment was assessed using a hypergeometric test with the Homo sapiens annotation (org.Hs.eg.db) as the reference background. GO terms with an adjusted p-value (Benjamini-Hochberg method) ≤ 0.05 were considered significantly enriched.

### Generating stable knockout and overexpression cell lines

CRISPR-mediated knockout of *SLC43A1* was achieved by co-transfecting SW620, RKO and 1A9 cells with CRISPR knockout and homology-directed repair plasmids (obtained from SantaCruz Biotechnology, Dallas, TX) using Lipofectamine 2000 (Invitrogen) and subsequent selection with puromycin (0.5mg/mL). Single-cell knockout clones were subsequently isolated, and protein levels were verified by western blot analysis. Overexpression of *SLC43A1* was achieved by transfecting SW620, RKO, and 1A9 OX80 cells with a pcDNA3.1-C-(k)DYK vector containing the *SLC43A1* ORF sequence or empty vector (obtained from Genscript, Piscataway, NJ) using Lipofectamine 2000 and subsequent selection with G418 (1mg/mL). Single cell overexpressing clones were subsequently isolated, and protein levels were verified by western blot analysis. Stable knockout and overexpressing cell lines were maintained in puromycin or G418, respectively.

### Western blot analysis

Whole cell lysates were extracted from cell pellets with RIPA buffer (50 mM Tris – HCl [pH 7.4], 1% Triton X-100, 10% glycerol, 0.1% SDS, 2 mM EDTA, and 0.5% deoxycholate, 50 mM NaCl) containing protease inhibitor cocktail (cat# 5871, Cell Signaling Technology, Danvers, MA). Following sonication, samples were centrifuged at 5000 x g for 10 minutes and soluble proteins in the supernatant were collected. Protein concentrations were determined using Bio-Rad Protein Assay Dye Reagent (Bio-Rad, Hercules, CA) according to the manufacturer’s instructions. Approximately 50-75 μg of protein lysates were analyzed by SDS-PAGE and transferred to nitrocellulose membranes. The membranes were probed with antibodies to detect LAT3 (Millipore-Sigma, #HPA018813-100UL), LAT1 (Cell Signaling #5347), LAT2 (Millipore-Sigma #HPA003462), LAT4 (Millipore-Sigma #HPA021564), phosphorylated γ-H2AX (Cell Signaling #2577), total γ-H2AX (Cell Signaling #2578), or beta-actin (generated in mouse cat# 3700, or rabbit cat# 4970, Cell Signaling) and mouse anti-alpha-tubulin (Sigma #T6074) as loading controls. After incubation with primary antibodies, membranes were incubated with fluorescently tagged IRDye secondary antibodies (LI-COR, Lincoln, NE, USA), and bands were visualized using the Odyssey CLx imaging system (LI-COR).

### Cytotoxicity assay

Cells were seeded in opaque white, 96-well plates at a density of 5,000 cells/well and allowed to adhere overnight. Cells were then treated with increasing concentrations of the desired compound and incubated for 72 h. Percentage of cell viability relative to untreated cells was determined using Cell TiterGlo (Promega, Madison, WI) according to the manufacturer’s instructions and reported as average of triplicate wells. The concentration at which 50% of cell growth was inhibited (GI50) was determined using GraphPad Prism. When indicated, cells were treated with pan-LAT inhibitor BCH or treated with platinum agents in the presence of excess amino acids using MEM Amino Acids Solution (ThermoFisher).

### Immunohistochemistry Analysis

Cells were grown under normal growth conditions to about 90% confluency, lifted, and collected in serum-free media. Cells were pelleted at 1000 x g for 5 minutes, clotted with thrombin/fibrinogen, 10% NBF fixed, and paraffin embedded. Sections were stained on the Bond RX autostainer (Leica Biosystems, Deer Park, IL) with LAT3 antibody (1:50 dilution) using EDTA retrieval buffer and nuclei counterstained with hematoxylin. Slides were scanned using the Aperio AT2 (Leica Biosystems) at 40X magnification.

### Colony formation assay

Cells were seeded at 1,000 cells/well in triplicate wells per condition and allowed to adhere overnight. SW620 cells were treated 3 times over the course of 12 days with 0-0.24µM oxaliplatin. RKO cells were treated twice over the course of 8 days with 0-0.5µM oxaliplatin. Colonies were stained with crystal violet at experimental endpoint, and whole well scans were obtained using ECHO Revolution Microscope. Colonies were counted using ImageJ, and survival fraction represents average number of colonies formed per condition, normalized to plating efficiency per cell line.

### qPCR analysis

RNA was extracted from cell pellets using Qiashredders and RNeasy Mini Kit (74104). cDNA was generated according to manufacturer’s instructions using iScript^TM^ cDNA Synthesis Kit (Bio-Rad). RT-qPCR analysis was performed using Bio-Rad CFX96 Real Time System with Taqman Fast Advanced Master Mix (4444557) and human Taqman primers (Thermo Fisher Scientific) as follows: SLC43A1 (Hs00992327_m1), SLC7A5 (Hs01001189_m1), SLC7A8 (Hs00794796_m1), SLC43A2 (Hs01018693_m1), SLC22A1 (Hs00427554_m1), SLC22A2 (Hs00533907_m1), SLC22A3 (Hs00222691_m1), SLC47A1 (Hs00217320_m1), SLC31A1 (Hs00977266_g1) and GAPDH (Hs_99999905_m1).

### Cellular uptake of Pt

Cells were seeded at 6.0-6.5 x 10^5^ cells/mL on 6-well plates and allowed to adhere overnight. Cells were either left untreated as a negative control or treated with 1 µM oxaliplatin or 1 µM cisplatin for 3 and 6 hours. Cells were washed once with ice-cold PBS, collected by cell scraper into ice-cold PBS, and centrifuged at 1,500 rpm for 15 minutes. Supernatant was discarded and cell pellet frozen at-80°C. Intracellular platinum levels were determined by inductively coupled plasma mass spectrometry (ICP-MS). Briefly, cell pellets were suspended in 100 µL 70% trace metal grade nitric acid (Millipore-Sigma, PN#NX0407) and heated to 90 °C for two hours. Following, 50 µL of trace metal grade hydrogen peroxide (Millipore-Sigma, PN#95321) was added to the pellet suspension, where heating was then resumed for an additional hour. Post heating, the pellet solution was diluted to 1 mL with water and elemental iridium (Inorganic Ventures, Christianburg, VA) for internal standard purposes. Samples were then injected via peristaltic pump into a Thermo iCAPQ ICP-MS (ThermoFisher Scientific) where data were collected, specifying quantitation of ^194^Pt, ^195^Pt and ^193^Ir. A standard curve was prepared using elemental platinum (Inorganic Ventures, Christianburg, VA) and iridium internal standard and concentrations determined via the ratio of ^194^Pt to ^193^Ir. ^194^Pt levels were normalized to average cell number per well determined at the time of cell harvesting. In experiments with excess amino acids, cells were plated and treated with oxaliplatin in normal growth media supplemented with MEM Amino Acids Solution (ThermoFisher). For experiments conducted in the absence of amino acids, cells were seeded in normal growth media, washed 1X with dPBS, and treated with oxaliplatin in PBS with calcium and magnesium.

### Assessment of DNA damage

Cells were seeded in 10cm dishes (2-2.5×10^6^), allowed to adhere overnight, and treated with oxaliplatin for 24 hours. Cells were washed once with ice-cold PBS, scraped, and centrifuged at 2,000 rpm for 5 minutes. For analysis of DNA-platinum adducts, DNA was extracted from cell pellets using DNeasy Blood & Tissue Kit (Qiagen #69504) and concentrations measured using Denovix DS-11+ Spectrophotometer. Platinum adducts were measured using ICP-MS. Samples were prepared similarly to previous reports, with some modifications (46). Briefly, 25 uL of DNA extracts were combined with 50 uL 70% trace metal grade nitric acid, and heated for 1 hour at 65 °C, then an equal volume of 30% trace metal grade hydrogen peroxide was added for heating for 3 hours. Post heating, the samples were analyzed as described above for the cell pellet samples. Quantitated platinum was normalized to DNA concentration (ng/uL). For γ-H2AX analysis, cells were treated and collected under the same conditions and cell pellets were prepared for immunoblotting.

### Data availability

CRISPR screen data were deposited in the Gene Expression Omnibus database with accession number GSE294894.

### Statistical analyses

Unless noted otherwise, all statistical analyses including standard error of mean (SEM) and statistical significance were determined using GraphPad Prism 10 (GraphPad Software, Boston, MA). Data were considered statistically significant when p < 0.05 using an unpaired two-tailed Student’s t-test.

