## Supplementary files for "A whole genomic CRISPR-Cas9 screen identifies the amino acid transporter *SLC43A1* (LAT3) as a major determinant of oxaliplatin sensitivity in colorectal cancer cells"

**SUPPLEMENTARY DATA**

**Figure S1. Genome-wide CRISPR screens of CRC cell lines reveal SLC43A1 as top hit correlated with oxaliplatin sensitivity. A)** sgRNA counts from SW620 and RKO knockout screen in T20 untreated and T0 control conditions, sorted by common essential or non-essential genes (45). **B)** Overlap of significant negative hits across all activation screen (left) and knockout screen (right) conditions ($p<0.05$, fold change ≥ 2). **C)** Gene Ontology (GO) Biological Processes for the top 500 most significantly depleted genes from the CRISPR knockout screens were identified using a hypergeometric test (Benjamini-Hochberg adjusted p-value ≤ 0.05). The x-axis of the dot plot represents the Gene Ratio, which is the proportion of CRISPR hits found within a given GO Biological Process relative to the total number of genes in that category. The size of each dot corresponds to the total number of CRISPR hits associated with that GO term. **D)** SW620 knockout screen; $\log_{10}$ RRA scores in cells treated with 0.38μM (*left*) or 0.5μM OxPt (*right*) at Day 20 compared to control untreated cells at Day 0. **E)** RKO knockout screen; $\log_{10}$RRA scores in cells treated with 1.9μM OxPt at Day 20 compared to control untreated cells at Day 0. **F)** -$\log_{10}$ RRA scores representing depleted genes in SW620 and RKO knockout screens in T20 untreated cells compared to T0 control cells. Top 5 essential genes that were significantly altered are labeled, as well as SLC43A1. **G)** SW620 activation screen; $\log_{10}$ RRA scores in cells treated with 0.38μM (*left*) or 0.5μM OxPt (*right*) at Day 20 compared to control untreated cells at Day 0. **H)** RKO activation screen; $\log_{10}$ RRA scores in cells treated with 1.9μM at Day 20 compared to control untreated cells at Day 0. Top 10 enriched or depleted hits in each screen are colored blue or lilac, respectively; data was analyzed using MAGeCKFlute (46). **I)** Log fold change of SLC43A1 sgRNA read counts from SW620 (*left*) and RKO (*right)* cells treated with OxPt at Day 20 relative to respective controls from knockout (KO) and activation (A) screens ($p<0.001$).

**Figure S2. LAT3 knockout and overexpression do not affect sensitivity to other platinum agents. A)** SW620 and **B)** RKO LAT3KO, or **C)** SW620 and **D)** RKO LAT3OE cells were treated with cisplatin (0-20μM), carboplatin (0-100μM) or phenanthriplatin (0-20μM) for 72 hours and cell viability was measured by CellTiterGlo. Left panels show representative dose response of average cell viability from triplicate wells; bar graphs show average GI50 values from 3-5 independent experiments. All error bars represent SEM. Data are represented as mean ± SEM; *p<0.05, **p<0.01, ns = not significant.

**Figure S3. LAT1 knockout and overexpression does not affect sensitivity to oxaliplatin. A)** Log-fold change values of SLC7A5 gRNAs in SW620 CRISPR KO screen, from cells treated with 0.5μM OxPt at Day 20 versus untreated cells at Day 0 (T0_T20) or versus untreated cells at Day 20 (T20_T20); p<0.01. **B)** Immunoblot analysis of SW620 parental cells, LAT1 knockout clones 1 and 2, pcDNA empty vector control, and LAT1 overexpressing cells (C9) for LAT1 and β-actin protein expression. **C)** SW620 LAT1KO and **D)** LAT1OE cells were treated with oxaliplatin (0-20μM) for 72 hours and cell viability was measured by CellTiterGlo. Representative dose response curve shows average cell viability from triplicate wells; bar graphs show average GI50 from 3-4 independent experiments. **E)** SW620 LAT1KO cells were treated with 0-0.24μM OxPt for 10-12 days and colonies were stained with crystal violet. Left graph shows average survival fraction (%). Representative images show whole well scans of colonies stained with crystal violet at experimental endpoint. **F)** SW620 LAT1KO cells were treated with 1μM oxaliplatin for 3 or 6 hours, washed, and cell pellets were collected to measure intracellular platinum accumulation via ICP-MS. Average $^{194}$Pt values were normalized to cell number, and data represents average fold change relative to parental SW620 cells from 4 independent experiments. **G)** Immunoblot analysis of LAT1, LAT2, and LAT4 protein levels with α-tubulin and β-actin as loading controls. **H)** qPCR analysis of average LAT1, LAT2, and LAT4 mRNA expression, normalized to GAPDH housekeeping control, from 2 independent experiments performed in triplicate. All error bars represent SEM. Data are represented as mean ± SEM; *p<0.05, ns = not significant.

**Figure S4. Oxaliplatin resistant ovarian cancer cells are cross-resistant to cisplatin and carboplatin**. **A-C)** Cells were treated with cisplatin or carboplatin and cell viability was measured by CellTiterGlo after 72 hours. Representative dose response curves show average cell viability from triplicate wells. Bar graphs represent average GI50s**. D)**

Cells were treated with 1μM cisplatin or carboplatin for 6 hours, washed, and cell pellets were collected to measure intracellular platinum accumulation via ICP-MS. Average $^{194}Pt$ values were normalized to cell number, and data represents average fold change relative to parental 1A9 cells. All error bars represent SEM. Data are represented as mean ± SEM; n = 3 independent experiments performed in triplicate; *p<0.05, ***p<0.005, ****p<0.005, ns = not significant.

**Figure S5. LAT3 expression does not regulate other suggested transporters of oxaliplatin. A)** $Log_2$ fold change and corresponding RRA-score of sgRNAs corresponding to other SLC transporters that appeared in the top 25 positively selected genes in the SW620 CRISPR KO screen (0.38μM) and top 25 negatively selected genes in the SW620 CRISPRa screen (0.38μM) (SLC45A4, SLC5A2, SLC16A2, SLC25A10, SLC22A25, SLC43A2, SLC31A1), and other suggested transporters of oxaliplatin (SLC22A1, SLC22A2, SLC22A3, ATP7A, ATP7B, ABCC1, ABCC2, SLC47A1). **B)** qPCR analysis of SLC22A21 (OCT1), SLC22A3 (OCT3), SLC47A1 (MATE1) and SLC31A1 (CTR1) mRNA expression normalized to GAPDH in SW620 and RKO LAT3KO and LAT3OE clones. SLC22A2 was undetected in both cell lines. Data represents average fold change relative to respective controls ± SEM, and unless otherwise indicated are not statistically significant (n = 2 independent experiments performed in triplicate; *p<0.05).

FIGURE S1

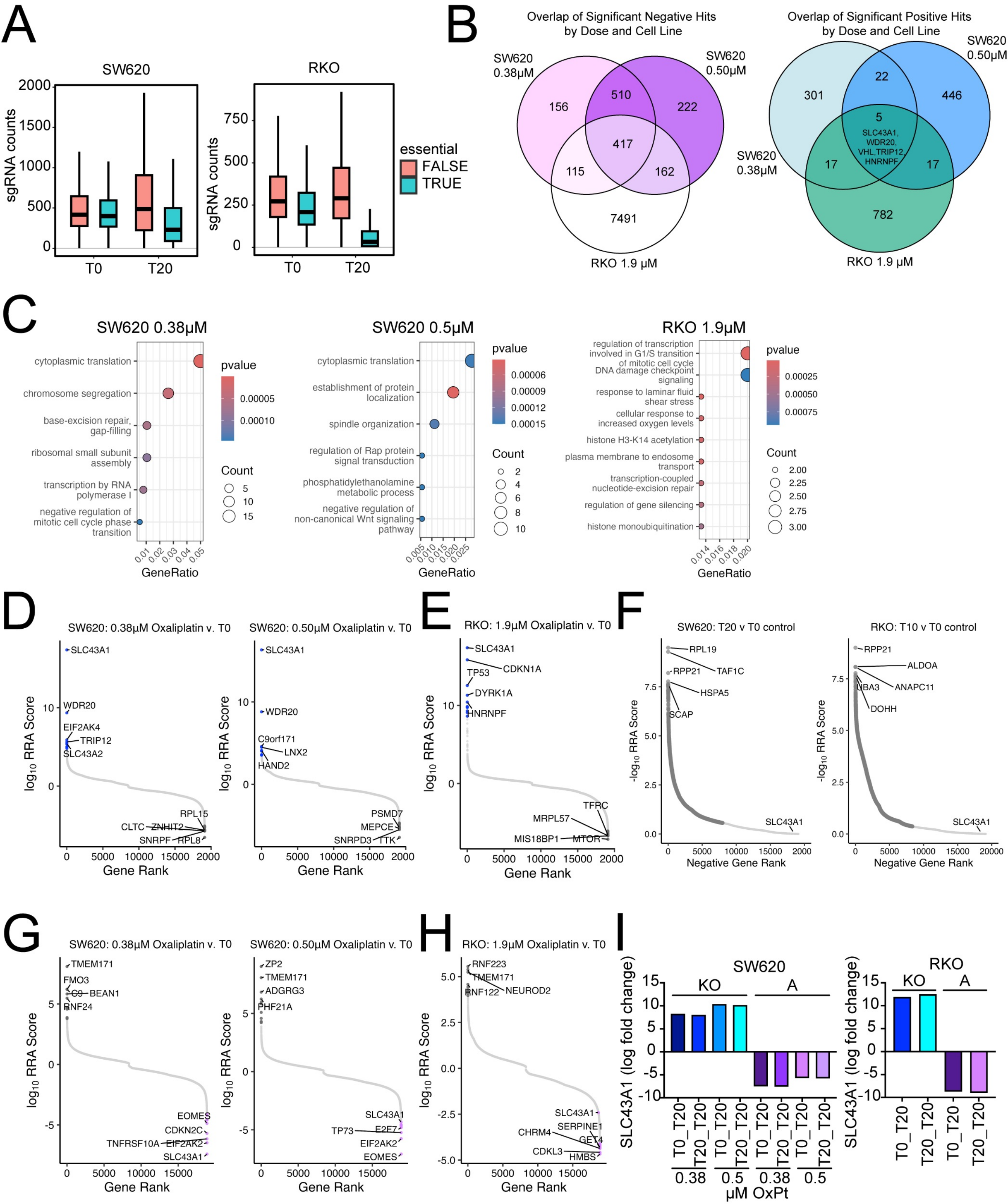
A
SW620
RKO
sgRNA counts
T0
T20
essential
FALSE
TRUE
B
Overlap of Significant Negative Hits by Dose and Cell Line
SW620 0.38μM
SW620 0.50μM
RKO 1.9 μM
156
510
222
417
115
162
7491
Overlap of Significant Positive Hits by Dose and Cell Line
301
22
446
5
SLC43A1, WDR20, VHL,TRIP12, HNRNPF
17
17
782
C
SW620 0.38μM
cytoplasmic translation
chromosome segregation
base-excision repair, gap-filling
ribosomal small subunit assembly
transcription by RNA polymerase I
negative regulation of mitotic cell cycle phase transition
pvalue
Count
GeneRatio
SW620 0.5μM
cytoplasmic translation
establishment of protein localization
spindle organization
regulation of Rap protein signal transduction
phosphatidylethanolamine metabolic process
negative regulation of non-canonical Wnt signaling pathway
RKO 1.9μM
regulation of transcription involved in G1/S transition of mitotic cell cycle
DNA damage checkpoint signaling
response to laminar fluid shear stress
cellular response to increased oxygen levels
histone H3-K14 acetylation
plasma membrane to endosome transport
transcription-coupled nucleotide-excision repair
regulation of gene silencing
histone monoubiquitination
D
SW620: 0.38μM Oxaliplatin v. T0
SW620: 0.50μM Oxaliplatin v. T0
log10 RRA Score
Gene Rank
SLC43A1
WDR20
EIF2AK4
TRIP12
SLC43A2
RPL15
CLTC
ZNHIT2
SNRPF
RPL8
C9orf171
LNX2
HAND2
PSMD7
MEPCE
SNRPD3
TTK
E
RKO: 1.9μM Oxaliplatin v. T0
CDKN1A
TP53
DYRK1A
HNRNPF
TFRC
MRPL57
MIS18BP1
MTOR
F
SW620: T20 v T0 control
RKO: T10 v T0 control
-log10 RRA Score
Negative Gene Rank
RPL19
RPP21
TAF1C
HSPA5
SCAP
ALDOA
UBA3
ANAPC11
DOHH
G
SW620: 0.38μM Oxaliplatin v. T0
SW620: 0.50μM Oxaliplatin v. T0
TMEM171
FMO3
C9
BEAN1
RNF24
EOMES
CDKN2C
TNFRSF10A
EIF2AK2
SLC43A1
ZP2
ADGRG3
PHF21A
TP73
E2F7
H
RKO: 1.9μM Oxaliplatin v. T0
RNF223
RNF122
NEUROD2
SERPINE1
CHRM4
GET4
CDKL3
HMBS
I
SW620
RKO
SLC43A1 (log fold change)
KO
A
T0_T20
T20_T20
0.38
0.5
μM OxPt

FIGURE S2

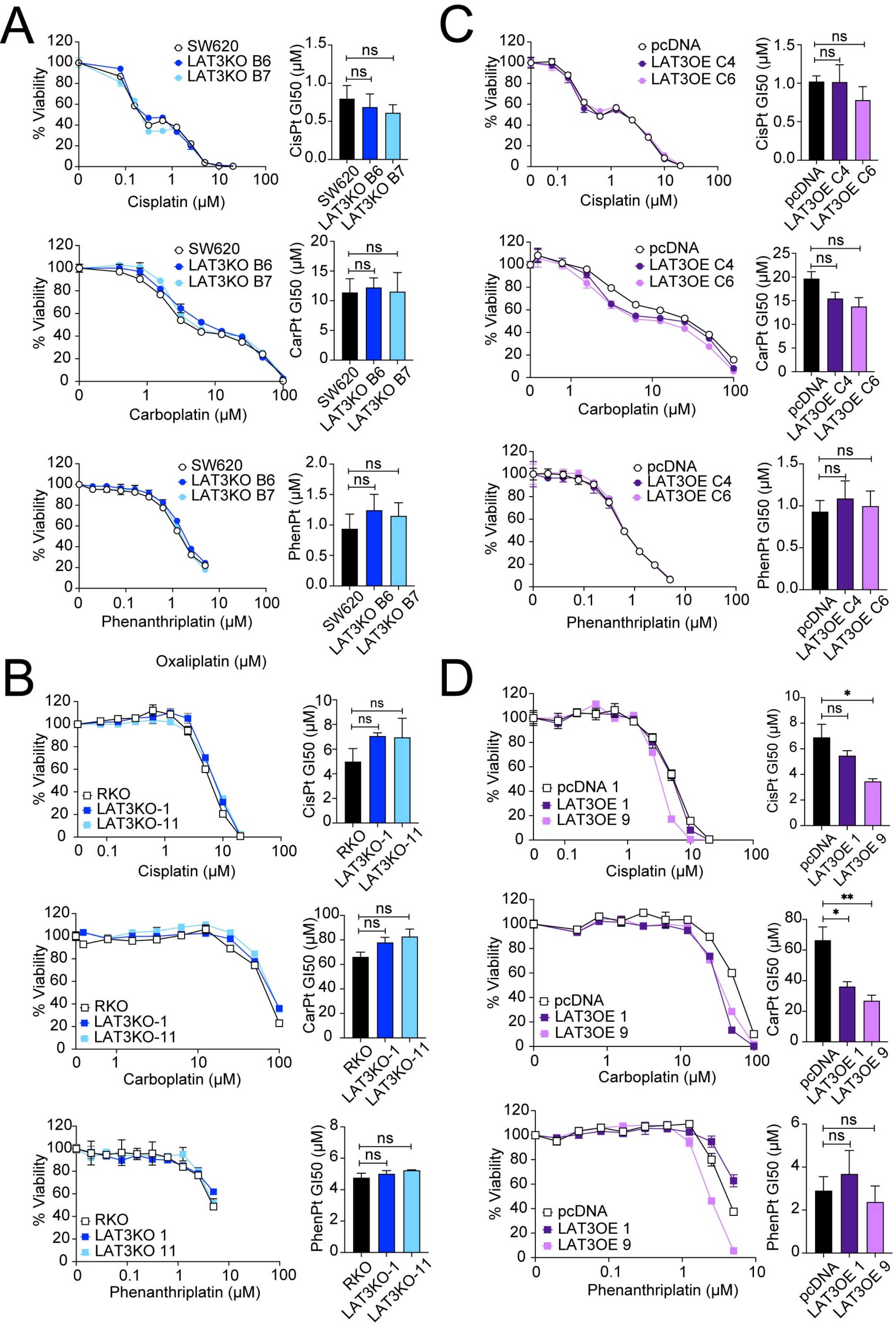
A
% Viability
SW620
LAT3KO B6
LAT3KO B7
Cisplatin (µM)
CisPt GI50 (µM)
ns
ns
Carboplatin (µM)
CarPt GI50 (µM)
Phenanthriplatin (µM)
PhenPt (µM)
Oxaliplatin (µM)
B
RKO
LAT3KO-1
LAT3KO-11
LAT3KO 1
LAT3KO 11
Cisplatin (µM)
CisPt GI50 (µM)
Carboplatin (µM)
CarPt GI50 (µM)
Phenanthriplatin (µM)
PhenPt GI50 (µM)
C
pcDNA
LAT3OE C4
LAT3OE C6
Cisplatin (µM)
CisPt GI50 (µM)
Carboplatin (µM)
CarPt GI50 (µM)
Phenanthriplatin (µM)
PhenPt GI50 (µM)
D
pcDNA 1
pcDNA
LAT3OE 1
LAT3OE 9
Cisplatin (µM)
CisPt GI50 (µM)
*
Carboplatin (µM)
CarPt GI50 (µM)
**
Phenanthriplatin (µM)
PhenPt GI50 (µM)

FIGURE S3

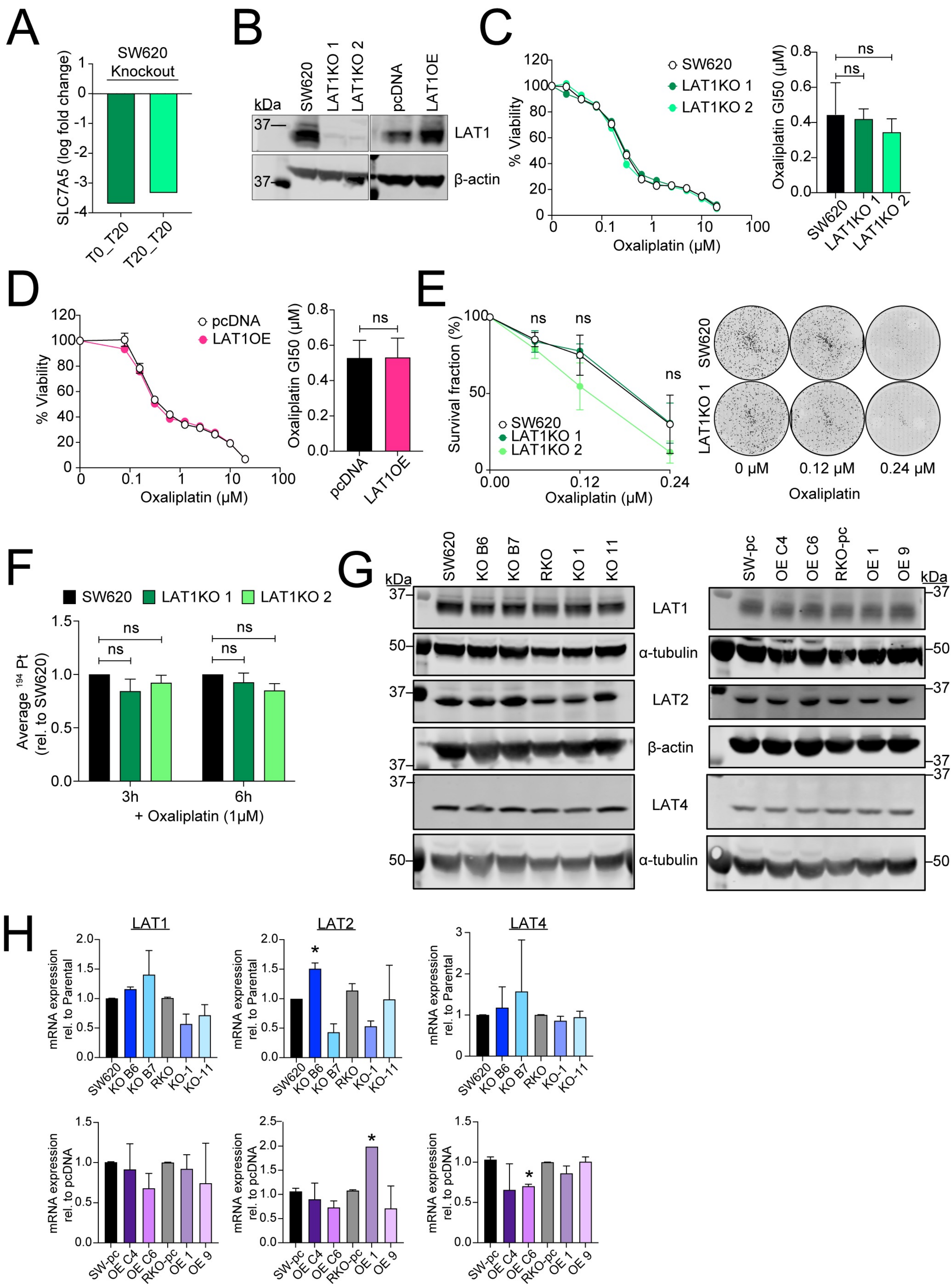

A
SW620
Knockout
SLC7A5 (log fold change)
T0_T20
T20_T20
B
kDa
SW620
LAT1KO 1
LAT1KO 2
pcDNA
LAT1OE
LAT1
β-actin
C
% Viability
Oxaliplatin (μM)
SW620
LAT1KO 1
LAT1KO 2
Oxaliplatin GI50 (μM)
ns
D
pcDNA
LAT1OE
E
Survival fraction (%)
0 μM
0.12 μM
0.24 μM
Oxaliplatin
F
Average 194 Pt (rel. to SW620)
3h
6h
+ Oxaliplatin (1μM)
G
KO B6
KO B7
RKO
KO 1
KO 11
SW-pc
OE C4
OE C6
RKO-pc
OE 1
OE 9
α-tubulin
LAT2
LAT4
H
mRNA expression rel. to Parental
mRNA expression rel. to pcDNA
KO-1
KO-11

FIGURE S4

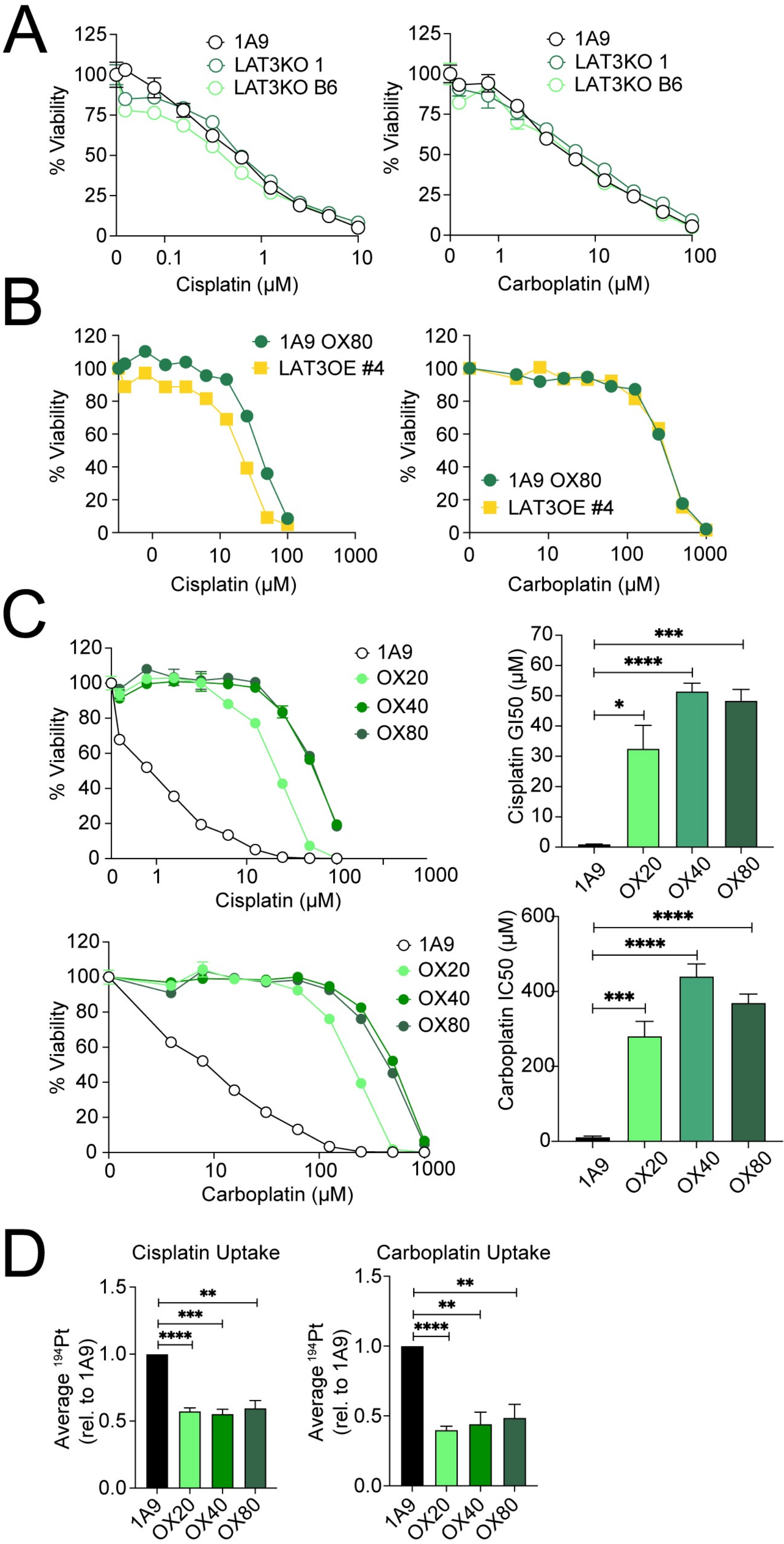

A
1A9
LAT3KO 1
LAT3KO B6
% Viability
Cisplatin (µM)
Carboplatin (µM)
B
1A9 OX80
LAT3OE #4
C
OX20
OX40
OX80
Cisplatin GI50 (µM)
Carboplatin IC50 (µM)
D
Cisplatin Uptake
Carboplatin Uptake
Average $^{194}$Pt (rel. to 1A9)

FIGURE S5

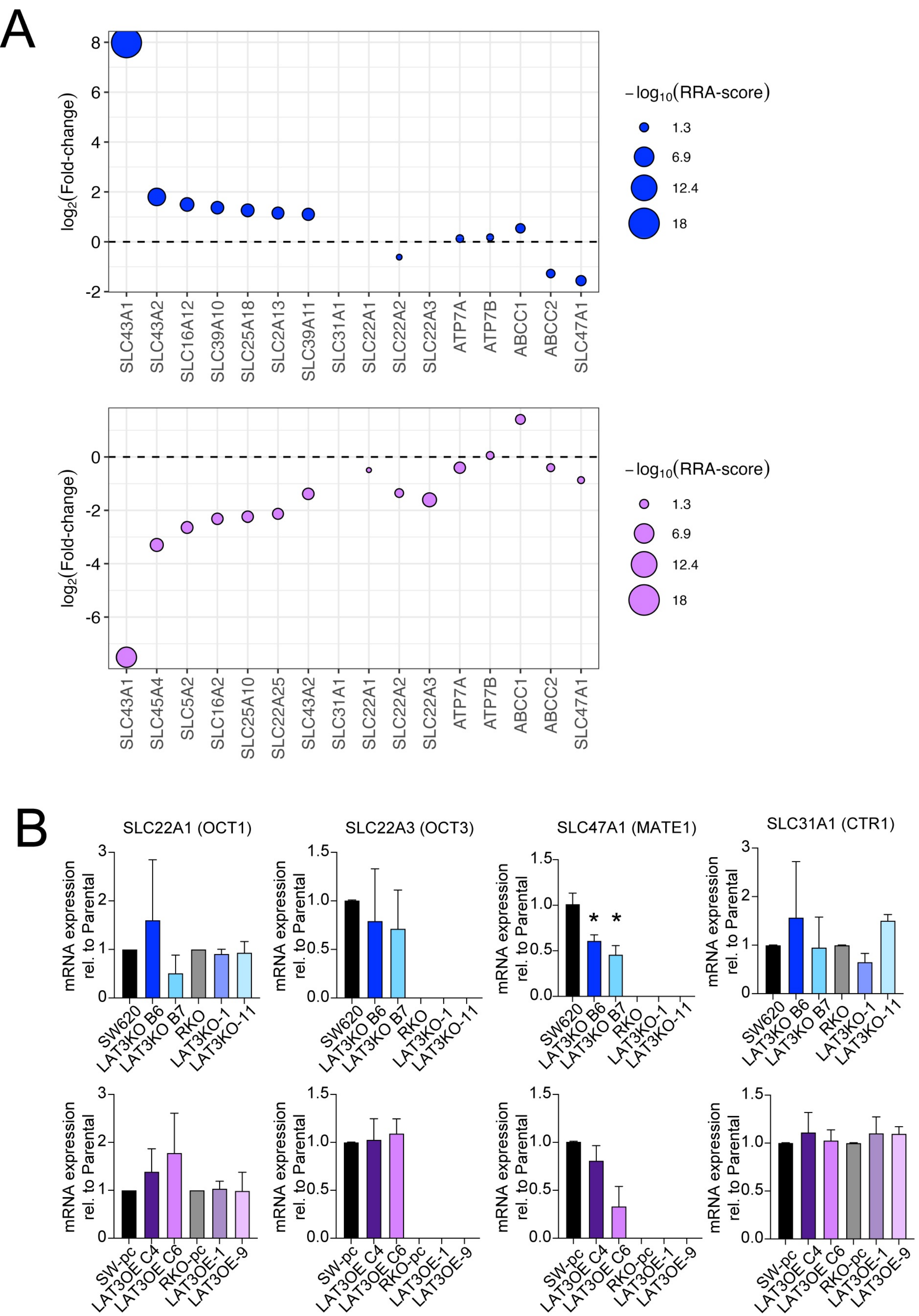
A
log2(Fold-change)
8
6
4
2
0
-2
−log10(RRA-score)
1.3
6.9
12.4
18
SLC43A1
SLC43A2
SLC16A12
SLC39A10
SLC25A18
SLC2A13
SLC39A11
SLC31A1
SLC22A1
SLC22A2
SLC22A3
ATP7A
ATP7B
ABCC1
ABCC2
SLC47A1
log2(Fold-change)
0
-2
-4
-6
−log10(RRA-score)
1.3
6.9
12.4
18
SLC43A1
SLC45A4
SLC5A2
SLC16A2
SLC25A10
SLC22A25
SLC43A2
SLC31A1
SLC22A1
SLC22A2
SLC22A3
ATP7A
ATP7B
ABCC1
ABCC2
SLC47A1
B
SLC22A1 (OCT1)
SLC22A3 (OCT3)
SLC47A1 (MATE1)
SLC31A1 (CTR1)
mRNA expression rel. to Parental
SW620
LAT3KO B6
LAT3KO B7
RKO
LAT3KO-1
LAT3KO-11
*
*
SW-pc
LAT3OE C4
LAT3OE C6
RKO-pc
LAT3OE-1
LAT3OE-9
